# Divergent effects of target protein stabilisation versus overproduction on PROTAC activity

**DOI:** 10.64898/2026.04.15.718695

**Authors:** Evelina Gudauskaitė, Brianda Hernández Morán, Gillian CA Taylor, Andrew J Wood

## Abstract

Protein degrader drugs such as PROTACs are being advanced as therapeutics targeted against oncogenic proteins. During tumorigenesis, oncogenic proteins can become constitutively activated via mechanisms including gene amplification, which increases protein production, and point mutations, which can extend protein half-life. Few experimental studies have addressed how disease-associated changes in target protein homeostasis influence PROTAC activity. We developed orthogonal methods to increase production or enhance stability of *β*-catenin, an important oncoprotein and target for degrader therapeutics, and used the dTAG system to evaluate the consequences for PROTAC activity. Stabilising oncogenic missense mutations increased protein expression up to 5-fold but do not alter the PROTAC-imposed minimal steady-state level. In contrast, transcriptional upregulation increases both pre- and post-treatment target levels, revealing a synthesis-dependent ceiling on achievable depletion. Our results highlight distinct constraints on PROTAC activity arising from different mechanisms of oncogene activation, with potential implications for preclinical modelling, drug resistance and personalised medicine.

## Introduction

The level of cellular protein expression is determined by the balance between rates of production and clearance. Protein degrader drugs such as PROTACs and molecular glues can induce rapid reductions in protein expression by inducing proximity between a target and an E3 ubiquitin ligase (Békés *et al*, 2022; Winter *et al*, 2015; Zengerle *et al*, 2015; Sakamoto *et al*, 2001). This leads to the formation of ubiquitin chains on the target, followed by rapid degradation via the proteasome. In this manner, degraders are thought to override endogenous protein stability pathways to reduce protein levels by shortening protein half-life.

At the time of writing, over 25 PROTACs have entered clinic trials, with many more in the preclinical pipeline. In many cases these molecules target oncogenic proteins, which become constitutively activated in particular subtypes of cancer (Hinterndorfer *et al*, 2025). Examples include the androgen receptor in prostate cancer, estrogen receptor in breast cancer, and KRAS in lung and pancreatic cancer. Preclinical programmes targeting other high value oncoproteins are being pursued by numerous groups globally.

During cancer initiation and progression, oncogenes can undergo activation via several distinct genetic mechanisms. Broadly speaking, these mechanisms can be distinguished based on whether they primarily affect the functional state, or the expression level, of the oncoprotein. For example, KRAS acts as an on/off switch for cellular growth by cycling between a GTP bound and unbound state; oncogenic mutations at the KRAS N-terminus (G12C/D/V) lock the protein in the active GTP-bound state (Uniyal *et al*, 2025).

Many other oncogenes are activated via mechanisms that increase the level of protein expression. Examples include cMYC, EGFR and SOX2, where gene loci are commonly amplified in cancer genomes, leading to increased rates of mRNA and protein production (Lockwood *et al*, 2008; Verhaak *et al*, 2019; Bass *et al*, 2009). In other cases such as ERG and NRF2, mutations disrupt interactions between degrons and E3 ubiquitin ligase receptors, which increases protein levels primarily via extending protein half-life rather than increasing rates of synthesis (Mészáros *et al*, 2017; Liu *et al*, 2021; Taguchi & Yamamoto, 2017). In other cases (e.g. MYCN), both gene amplification and point mutations have been reported in the same tumour (Williams *et al*, 2015; Otto *et al*, 2009). Thus, when oncoprotein dosage is upregulated in disease, the extent to which this is driven by changes in synthesis or degradation is a potential source of both inter- and intra-tumour heterogeneity.

Despite huge interest in protein degrader drugs, few empirical studies have tested the consequence of clinically relevant mutations which act by changing protein homeostasis on PROTAC activity. Intuitively, it should be easier to shorten the half-life of a protein that is naturally long-lived compared to a naturally unstable molecule, and this principle has been expressed in mathematical models (Bartlett and Gilbert 2022, Vetma *et al* 2024). A recent study further showed that fusion of a HiBit tag to the CRAF protein reduced its stability, and also reduced the Degradation maximum (D_max_) defined as the maximum percentage reduction in target protein levels achievable via PROTAC treatment (Vetma *et al*, 2024). Conversely, weak PROTAC activity has been observed against target proteins expressed from transiently transfected plasmids (Riching *et al*, 2018), which are typically present at high copy number. Despite these insights, it remains unclear whether activation of an individual PROTAC target protein at the level of production or stability could be a determining factor in the outcome of degrader drug treatment.

In this paper we focus on *CTNNB1*, which encodes *β*-catenin, a transcriptional coactivator and intracellular transducer of the canonical Wnt signalling pathway, a focus of current degrader drug development efforts (Simonetta *et al*, 2019; Gowans *et al*, 2024; Boudreau *et al*, 2025) and a dependency in many solid tumours. *β*-catenin is most frequently activated at the level of protein stability, usually via missense mutations within an N-terminal degron motif (Krishna *et al*, 2026; Morin *et al*, 1997), or via loss of the destruction complex subunit APC (Morin *et al*, 1997; Korinek *et al*, 1997). Less frequently, mutant alleles undergo copy number gain to increase the rate of *β*-catenin production (Rebouissou *et al*, 2016; Krishna *et al*, 2026). Here, we use degron tagging in a controlled cellular system to empirically determine the consequences of enhancing the stability versus increasing the production of *β*-catenin protein on degrader drug activity.

## Results

To develop a model system to study the degradation of *β*-catenin proteins with different turnover kinetics, we first generated a tagged construct in which FKBP12^F36V^ and GFP were fused to the C-terminus of the wildtype *CTNNB1* open reading frame (Fig. 1A). FKBP12^F36V^ is a degron tag of 12KDa (‘dTAG’), that enables the rapid proteasomal degradation of fusion proteins using pre-validated PROTACs that recruit either the CRL2^CRBN^ (dTAG-13) or CRL2^VHL^ (dTAGv1) E3 ubiquitin ligase complex ((Nabet *et al*, 2018, 2020) Figure 1B), and GFP enables sensitive protein quantification at the single cell level. The tags were fused via glycine-rich linker peptides to minimise steric interference (Figure 1C) and the constructs were integrated into a landing pad in the genome of Flp-In 293 cells using Flp/FRT recombination (Figure 1A), where expression was driven by the CMV promoter (Figure 1D). In subsequent experiments, this system enabled comparison between variant proteins stably expressed from the same integration site, driven by the same transcriptional regulatory elements.

**Figure 1:**
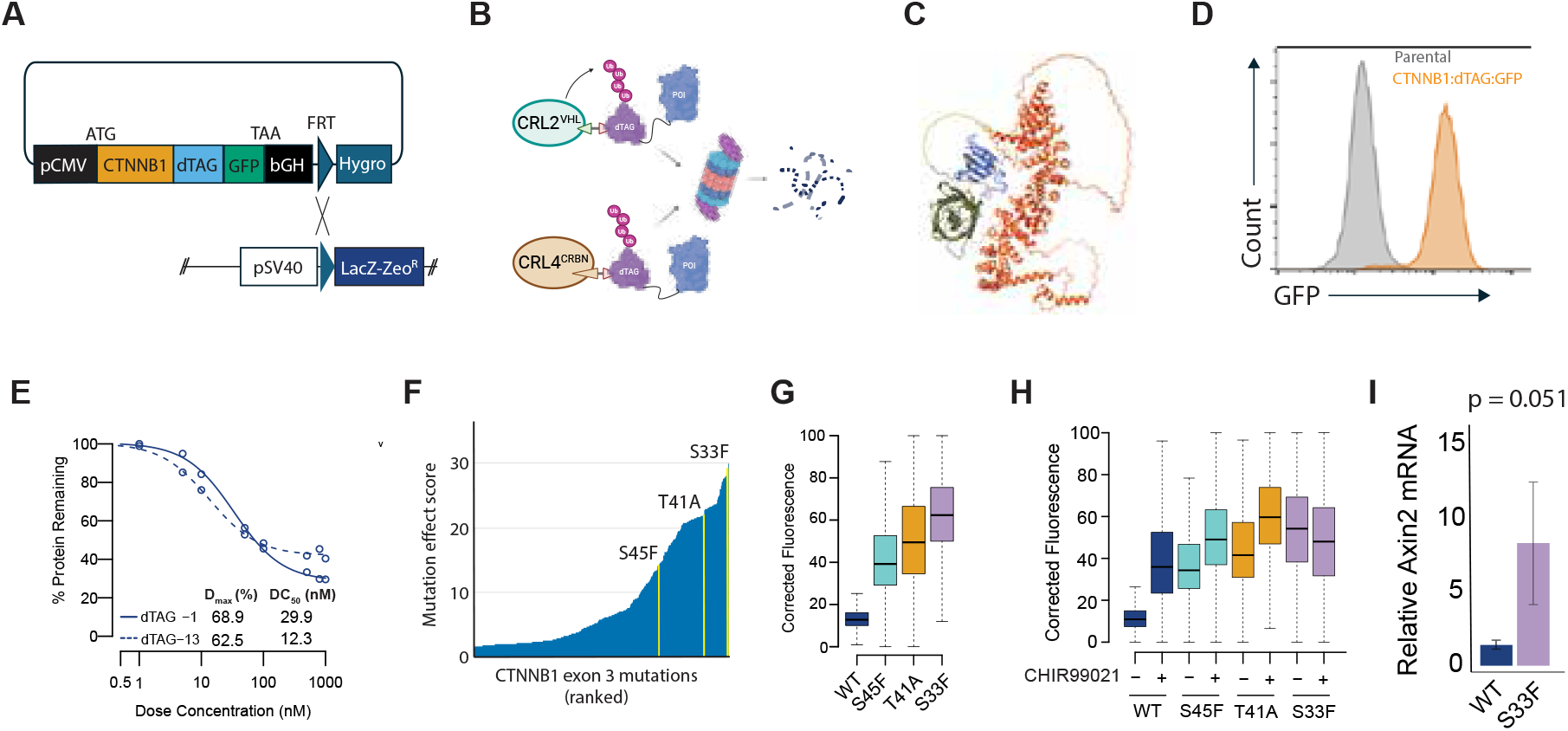
A dual degron tag reporter system to quantify CTNNB1 degradation kinetics. A. Schematic diagram of the CTNNB1:GFP:dTAG reporter construct. The entire pCDNA5 expression plasmid was integrated at a safe harbour locus in Flp-In 293 cells by co-transfection with a Flp recombinase expression vector (pOG44), then cells carrying stable integrations were selected using hygromycin. B. Schematic of the dTAG system. A hole modified FKBP12 (FKBP12^F36V^) tag is fused to the protein of interest. Two different PROTAC molecules then enable recruitment of either Cullin Ring E3 ligases CRL2^VHL^ (dTAGv1) or CRL4^CRBN^ (dTAG-13) to the tagged protein to facilitate poly-ubiquitination and destruction via the proteasome. C. Alphafold model of the CTNNB1 fusion protein, including FKBP12/dTAG (blue) and GFP (green). D. Measurement of steady-state GFP fluorescence in HEK293T cells stably expressing the CTNNB1 degron reporter from the Flp-In landing pad. E. Dose response curve showing degradation of the CTNNB1 degron reporter following 2 hour treatment with dTAGv1 or dTAG-13. Each point represents the median value from >1000 single cells, normalised to the median expression in vehicle treated cells. Data from a second independent experiment are shown in Figure S1A. F. Histogram showing the relative effects of all 342 possible missense mutations in the CTNNB1 native degron (amino acids 31 – 48), ranked according to the level that they activated a Tcf:GFP reporter protein (based on data from (Krishna *et al*, 2026)). Individual mutations used in this study are marked in in yellow. G. Boxplots show steady-state GFP expression levels in 293 cells stably expressing CTNNB1 degron reporters with different native degron mutations. In this and subsequent Figure panels, horizontal lines show the median fluorescence, boxes show the interquartile range and whiskers show the range. Data are derived from >1000 single cells, and are representative of measurements taken on at least two different days. H. Boxplots show the effect of inhibiting the destruction complex kinase GSK3b using CHIR99021 on steady-state CTNNB1 expression. This had a strong effect on WT protein but relatively weak effects on protein carrying native degron mutations. Data are representative of two independent experiments each measuring >1000 single cells per condition. I. Measurement of AXIN2 transcript levels by qRT-PCR shows that degron reporter proteins retain the ability to regulate endogenous target genes. p-value represents a one-tailed t-test. The ratio of Axin2 to *β*-Actin was measured in cells expressing the WT degron reporter, then the ratio in cells expressing the S33F mutant allele was expressed relative to this WT value.

Following antibiotic selection of recombinant cells, target protein degradation was quantified using flow cytometry over a 2 hour dose response to dTAG-13 or dTAGv1 (Figure 1E, Figure S1A). The two compounds elicited similar effects on target levels (Figure 1E, Figure S1A), with dTAGv1 being slightly more efficacious (Degradation maximum (D_max_) of 69% vs 58%) and dTAG-13 slightly more potent (Concentration required for 50% of maximum degradation (DC_50_) 13nM vs 34nM). Fusion of dTAG:GFP to the *β*-catenin C-terminus therefore provides a tractable system to induce and quantify PROTAC-mediated degradation.

Genetic variants within an endogenous degron spanning amino acid positions 32 – 45 of *β*-catenin (referred to hereafter as the ‘native’ degron’) are among the most frequent driver mutations in several human cancers ((Gao *et al*, 2017, Krishna *et al*, 2026). These mutations increase *β*-catenin protein dosage and downstream signalling by preventing recognition by the destruction complex, which includes protein kinases GSK3b and CK1a together with scaffolding proteins APC and AXIN1. When Wnt signalling is inactive during normal tissue development, this complex phosphorylates the native degron at positions S45, T41, S37 and S33, allowing recognition and ubiquitylation of *β*-catenin by the SCF^bTrCP^ E3 ubiquitin ligase, followed by proteasomal degradation (Clevers & Nusse, 2012). Mutations affecting these amino acids therefore extend protein half-life by preventing *β*-catenin destruction. Using a saturation genome editing assay, we recently showed that different missense mutations within the native degron exert distinct levels of effect on oncogenic signalling (Figure 1F), which influence the interaction of tumours with the immune system in hepatocellular carcinoma (Krishna *et al*, 2026).

To better understand the impact of oncogenic mutations that stabilise PROTAC targets, we selected 3 different variants (S45F, T41A, S33F) previously shown to induce *β*-catenin signalling at different levels: S45F activates relatively weakly, T41A activates to an intermediate degree, and S33F is a strong activator (Figure 1F, (Krishna *et al*, 2026)). These mutations were introduced into the *CTNNB1* coding sequence, and the resulting constructs were integrated at the genomic landing pad in HEK293T cells, as described above (Figure 1A). Each mutation increased the steady-state level of *β*-catenin protein expression to the expected degree: S45F increased GFP fluorescence by approximately 3-fold, T41A by 4-fold and S33F by 5-fold, compared to CTNNB1 lacking oncogenic mutation (Figure 1G). Expression differences between wildtype and mutant *β*-catenin proteins were greatly reduced following treatment with the GSK3*β* inhibitor CHIR99021 (Figure 1H), which prevents phosphorylation of serine and threonine residues within the degron. This demonstrates that the native *β*-catenin degron retains its regulatory function in the context of the dTAG:GFP fusion protein. Fusion proteins also showed the expected accumulation in the nucleus and cytoplasm in response to oncogenic mutations (Figure S2) and retained the ability to activate the *β*-catenin target gene *AXIN2* (Figure 1I). Fusion proteins therefore retain transcriptional regulatory function, consistent with a previous report (Ambrosi *et al*, 2022).

We next tested the consequence of oncogenic stabilising *CTNNB1* mutations on dTAG-13 and dTAGv1 activity using time course and dose response assays. Strikingly, the mutation-dependent differences in *β*-catenin expression that were evident before treatment (Figure 1G) were largely eliminated following 2 hour PROTAC exposure at doses above 500nM (Figure 2A, Figure S2B). Similarly, following treatment of cells expressing either wildtype or mutant CTNNB1 with 100nM PROTAC, absolute protein levels converged to essentially the same minimum value following prolonged exposure (Figure 2B, Figure S2C). Consequently, the mutation-dependent differences in *β*-catenin expression that were evident before treatment (Figure 1G) were not consistently observed at the 24 hour time point (Figure 2C, Figure S1D). Of note, when protein degradation was measured based on fractional reduction of signal (in line with common practice in the field), wildtype protein appeared to be degraded less effectively than protein with native degron mutations (Figure 2D, 2E, 2F). This was attributable to native degron mutations increasing protein levels before (Figure 1G) rather than after (Figure 2C, Figure S1D) treatment. In summary, native degron mutations enhance *β*-catenin steady-state expression before treatment, but PROTAC-mediated recruitment of either the CRL2^VHL^ (dTAGv1) or CRL4^CRBN^ (dTAG-13) E3 ligase complexes can push absolute protein levels to the same baseline minimum.

**Figure 2:**
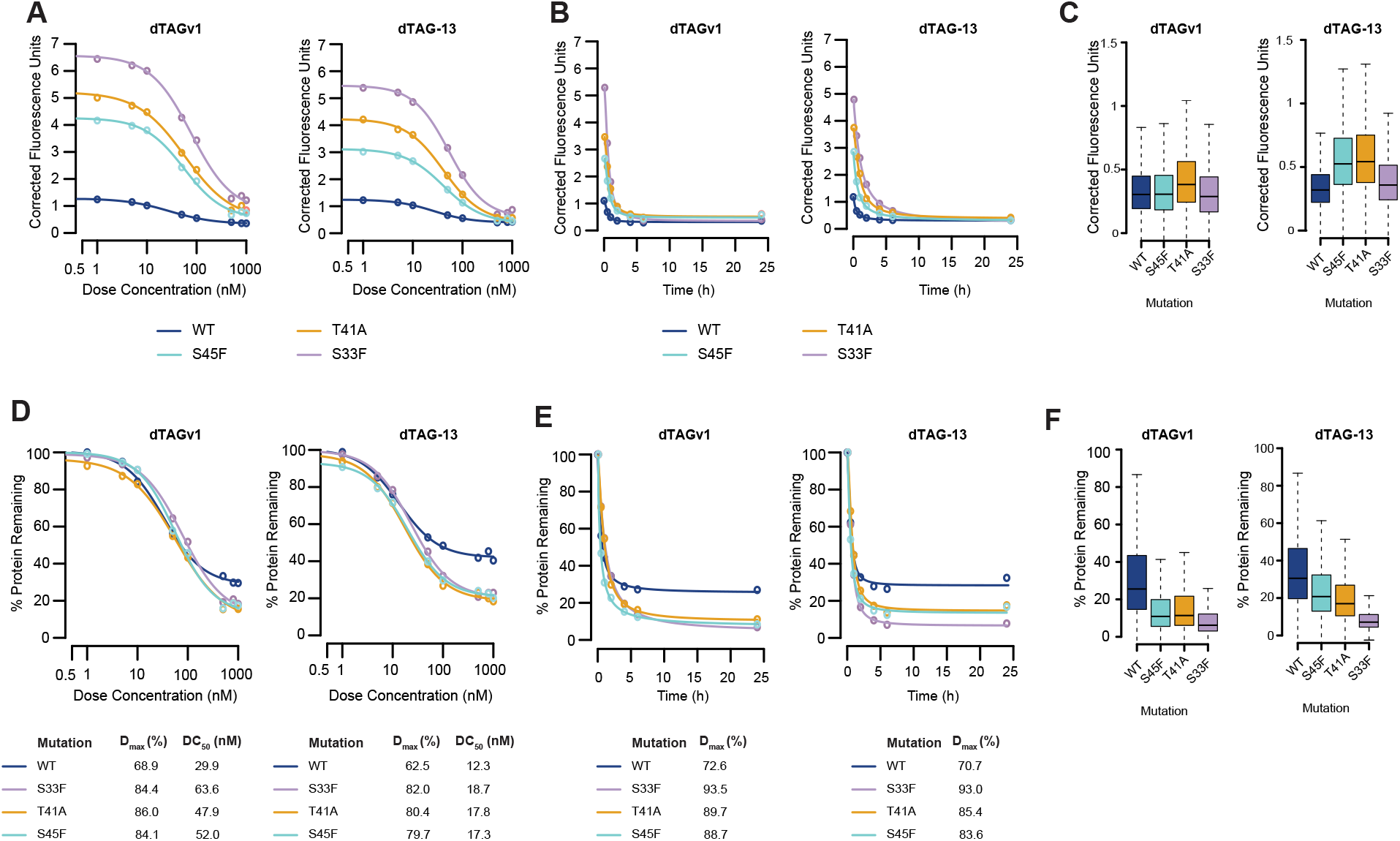
Stabilising CTNNB1 mutations differentially affect fractional versus absolute degradation. A. Dose response curves show the absolute level of GFP signal remaining following 2 hours of treatment with dTAGv1 (left) or dTAG-13 (right). Cell lines with either no mutation (WT) or specific missense mutations in the native degron are shown separately. Fluorescence units were corrected by subtracting the mean GFP signal from untreated non-transgenic 293 cells. Measurements were taken from >1000 single cells, and data from a second independent experiment are shown in Figure S1B. B. The level of corrected GFP signal is shown for proteins expressed from wildtype or native degron mutant transgenes over a 24-hour time course of treatment with dTAGv1 (left) or dTAG-13 (right) at 100nM, presented as described in panel A. Data from a second independent experiment are shown in Figure S1C. C. Box plots show the distribution of corrected cellular fluorescence values in >1000 single cells after 24-hour treatment with either dTAGv1 (left) or dTAG-13 (right). Horizontal lines show the median fluorescence, boxes show the interquartile range and whiskers show the range. Data from a second independent experiment are shown in Figure S1D. D. Dose response curves based on the same data shown in panel A, but fluorescence values are normalised to the level of corrected GFP signal in vehicle treated cells of the same genotype. The effect of exon 3 genotype on PROTAC D_max_ and DC_50_ values is shown below the plots. E. Time course data from panel B are presented with fluorescence expressed as a percentage of corrected GFP signal in vehicle treated cells of the same genotype. The effect of exon 3 genotype on PROTAC D_max_ and DC_50_ values is shown below the plots. F. Box plots show the same data as panel C, normalised to the level of corrected GFP signal in vehicle treated cells of the same genotype.

Although *β*-catenin is activated in cancer primarily via stabilising mutations, accumulating evidence suggests that regulation can also occur at the level of protein production (Rebouissou *et al*, 2016; Krishna *et al*, 2026; Chan *et al*, 2022). Across several wnt-dependent tumour types in the pan-cancer TCGA dataset, we found that gain or loss of the *CTNNB1* locus was associated with increased and decreased *CTNNB1* transcript levels, respectively (Figure 3A). Irrespective of copy number changes, levels of *CTNNB1* mRNA correlated moderately but consistently with transcript levels of downstream *β*-catenin target genes (Figure 3B). These data, combined with the knowledge that many other oncogenes are activated primarily at the level of transcriptional induction (Lockwood *et al*, 2008; Verhaak *et al*, 2019; Bass *et al*, 2009), prompted us to investigate whether upregulation of oncogene transcription could impose different constraints on PROTAC degradation profiles compared to changes in protein stability.

**Figure 3:**
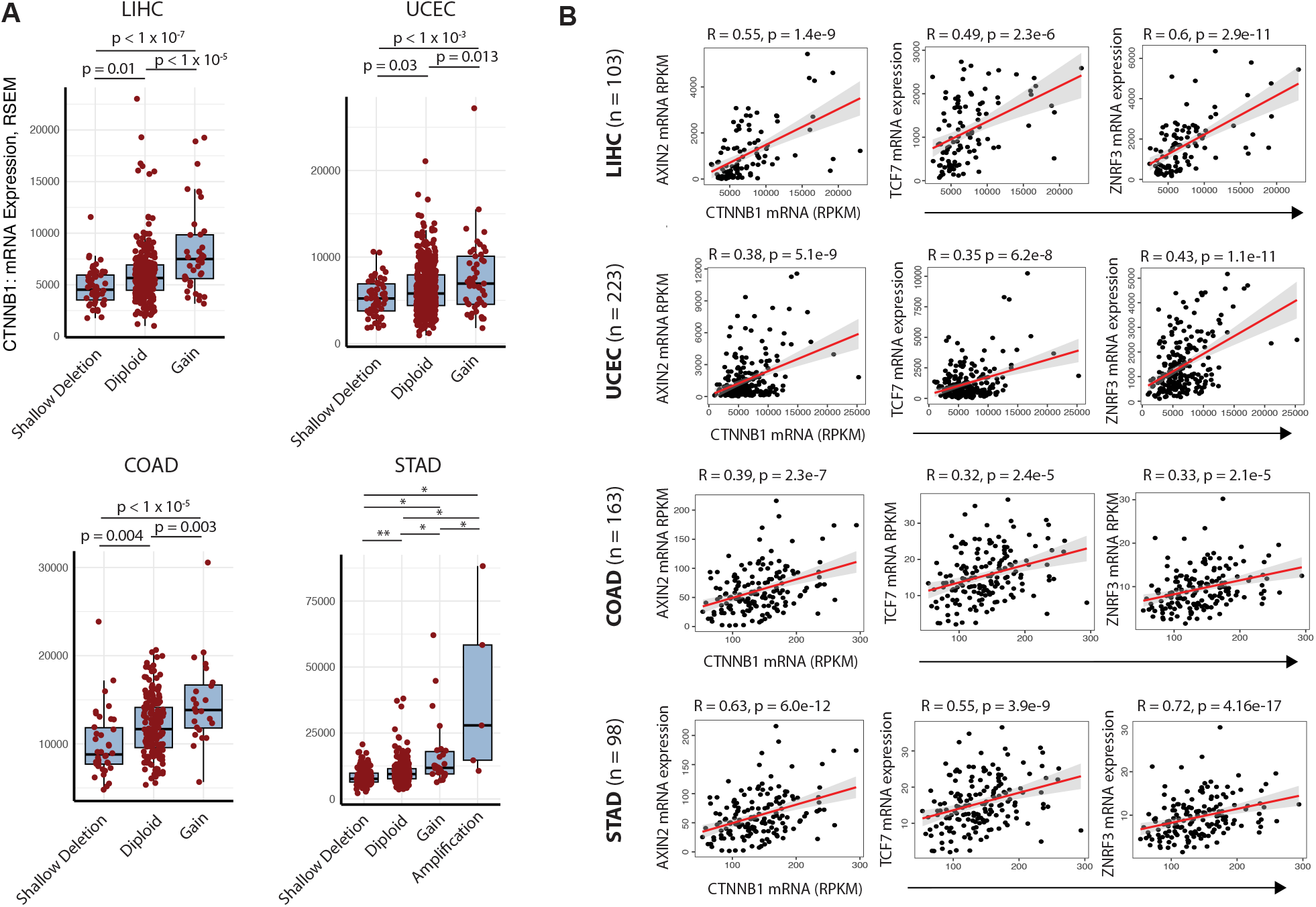
CTNNB1 mRNA levels correlate with target gene expression in Wnt-active tumours. A. Box and scatter plots show the distribution of CTNNB1 mRNA levels in samples with changes in CTNNB1 gene copy number, based on GISTIC 2.0 categories. All samples from LIHC (Hepatocellular Carcinoma), UCEC (Endometrial Carcinoma), COAD (Colorectal Adenocarcinoma) and STAD (Stomach Adenocarcinoma) cohorts from TCGA were analysed, but only GISTIC categories with 5 or more samples in any given tumour type are shown. Horizontal lines show median values, boxes show the interquartile range, whiskers show the range, and red points show individual samples. P values represent one-way ANOVA with Tukey’s post-hoc correction. For STAD, * denotes p < 1 x 10-7, ** denotes p < 0.05. B. Scatter plots show the relationship between mRNA levels of CTNNB1 and three downstream targets in the same four Wnt-dependent human tumour types detailed in panel A. To minimise the effects of non Wnt-active tumours within each class, only tumours with mutations in CTNNB1, APC or RNF43 are shown. R values are from Pearson correlations, linear regression lines are shown in red with the 95% confidence region shaded grey.

To address this, we modified our experimental system to enable transcriptional activation of *CTNNB1:dTAG:GFP*. Open reading frames encoding either WT, or S45F mutant *CTNNB1:dTAG:GFP* constructs were introduced into a landing pad in the genome of T-Rex 293 cells using Flp/FRT recombination (Figure 4A). T-Rex cells constitutively express the tet activator protein, and the landing pad promoter contains tet operator sequences that allow doxycycline inducible induction of transgene expression. Dox induction at 2.25*µ*M for 24 hours increased protein expression by a factor of 2.8-fold (WT) or 5.2-fold (S45F) over the basal state (Figure 4B), which was not increased further by longer or higher dosing.

**Figure 4:**
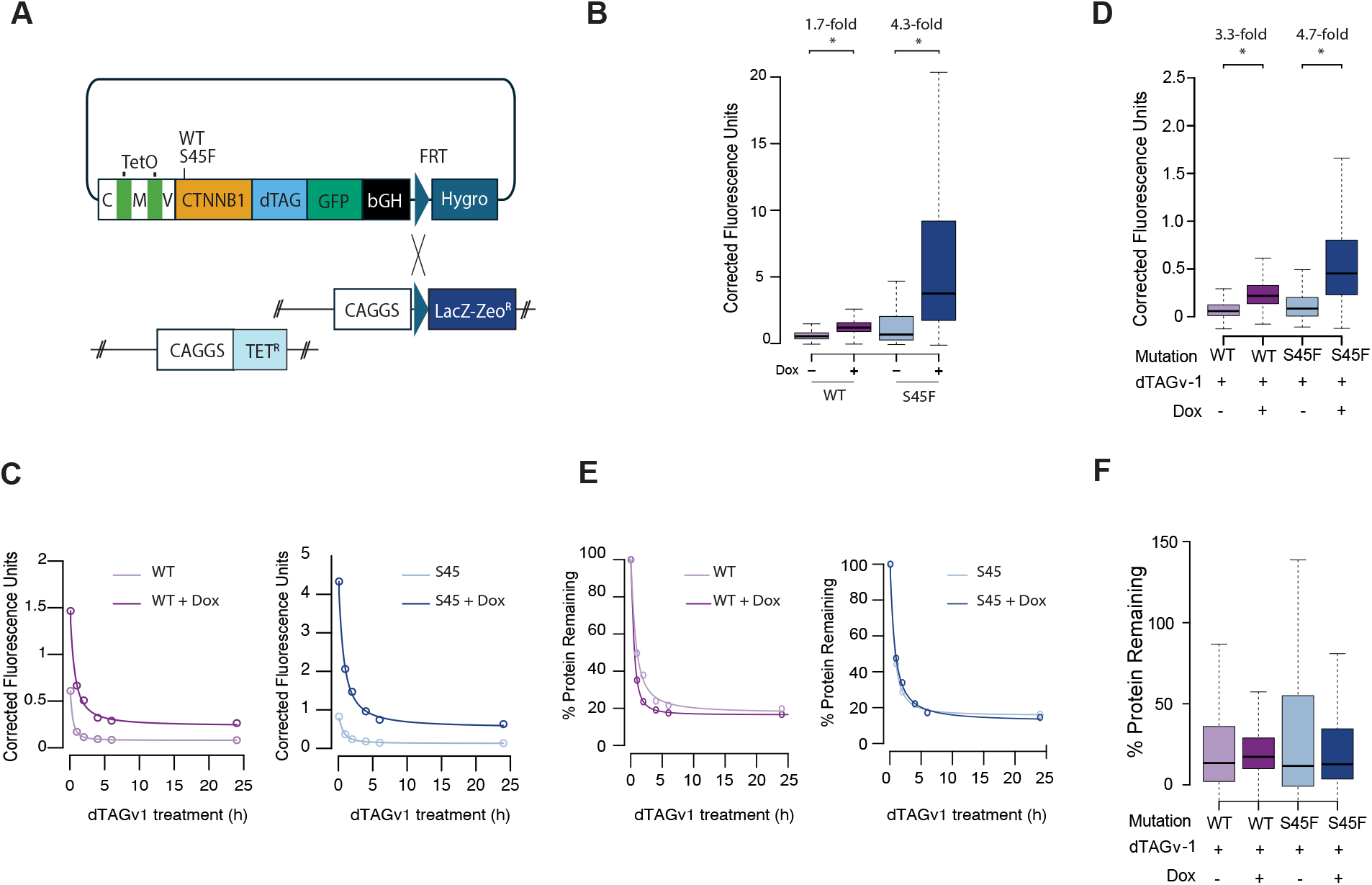
Transcriptional induction of CTNNB1 changes steady-state protein expression following PROTAC treatment. A. Schematic illustration showing the generation of 293 cells stably expressing CTNNB1 constructs transcribed from a doxycycline-inducible promoter (T-Rex). The entire plasmid was recombined into T-Rex cells already expressing the Tet activator protein (rtTA). B. In this and subsequent panels, boxplots show the change in CTNNB1 protein expression resulting from doxycycline induction of CTNNB1 transcription. Horizontal lines show median values of >1000 cells per condition, boxes show the interquartile range, whiskers show the range. Asterisk indicates statistically significant differences between conditions at p < 2.2 x 10-16 using a Mann Whitney U-test. C. Time course experiments show changes in absolute protein levels following dTAGv1 treatment at 100 nM. Each point represents mean background-corrected fluorescence levels from >1000 single cells. Data from a second independent experiment are shown in Figure S1E. D. Box plots show the distribution of corrected cellular fluorescence values in >1000 single cells after 24-hour treatment with dTAGv1 at 100nM. Asterisk indicates statistically significant differences between groups at p < 2.2 x 10-16 using a Mann Whitney U-test. E. Data from panel C are presented as % reduction in GFP signal compared to vehicle. Data from a second independent experiment are shown in Figure S1F. F. Box plots show the same data as panel D, normalised to the level of corrected GFP signal in vehicle treated cells of the same genotype.

Time course experiments were performed to quantify PROTAC activity in the presence or absence of prior transcriptional induction via 24 hour doxycycline treatment. A majority of *β*-catenin was degraded in both plus and minus doxycycline conditions (Figure 4C). Nonetheless, transcriptional induction increased absolute levels of target protein expression after 24 hours of PROTAC treatment (Figure 4D). In contrast to activation via native degron mutation (Figure 1G, Figure 2C), transcriptional induction was associated with elevated target protein dose both before (Figure 4B) and after (Figure 4D) PROTAC treatment, so the fractional degradation maximum did not change (Figure 4E, 4F). The results of transcriptional induction experiments were consistent for *CTNNB1* constructs with wildtype and S45F mutant native degrons (Figure 4B, C, D, E, F). Oncogene activation via transcriptional induction therefore has distinct effects on PROTAC activity compared to protein stabilisation via native degron inactivation.

## Disscusion

In this study, we investigated how distinct oncogenic activation mechanisms (altered protein stability versus transcriptional induction) shape the activity of proteolysis-targeting chimeras (PROTACs). Using *β*-catenin as a model oncogenic substrate, and FKBP12^F36V^ (dTAG) degraders as model PROTAC ligands (Nabet *et al*, 2018, 2020), we show that pretreatment proteostasis conditions fundamentally influence degradation outcomes, but in ways that differ depending on whether increased protein expression arises from degron inactivation or transcriptional induction (Figure 5). These findings provide a conceptual framework for anticipating degrader behaviour across genetically diverse tumour contexts.

**Figure 5:**
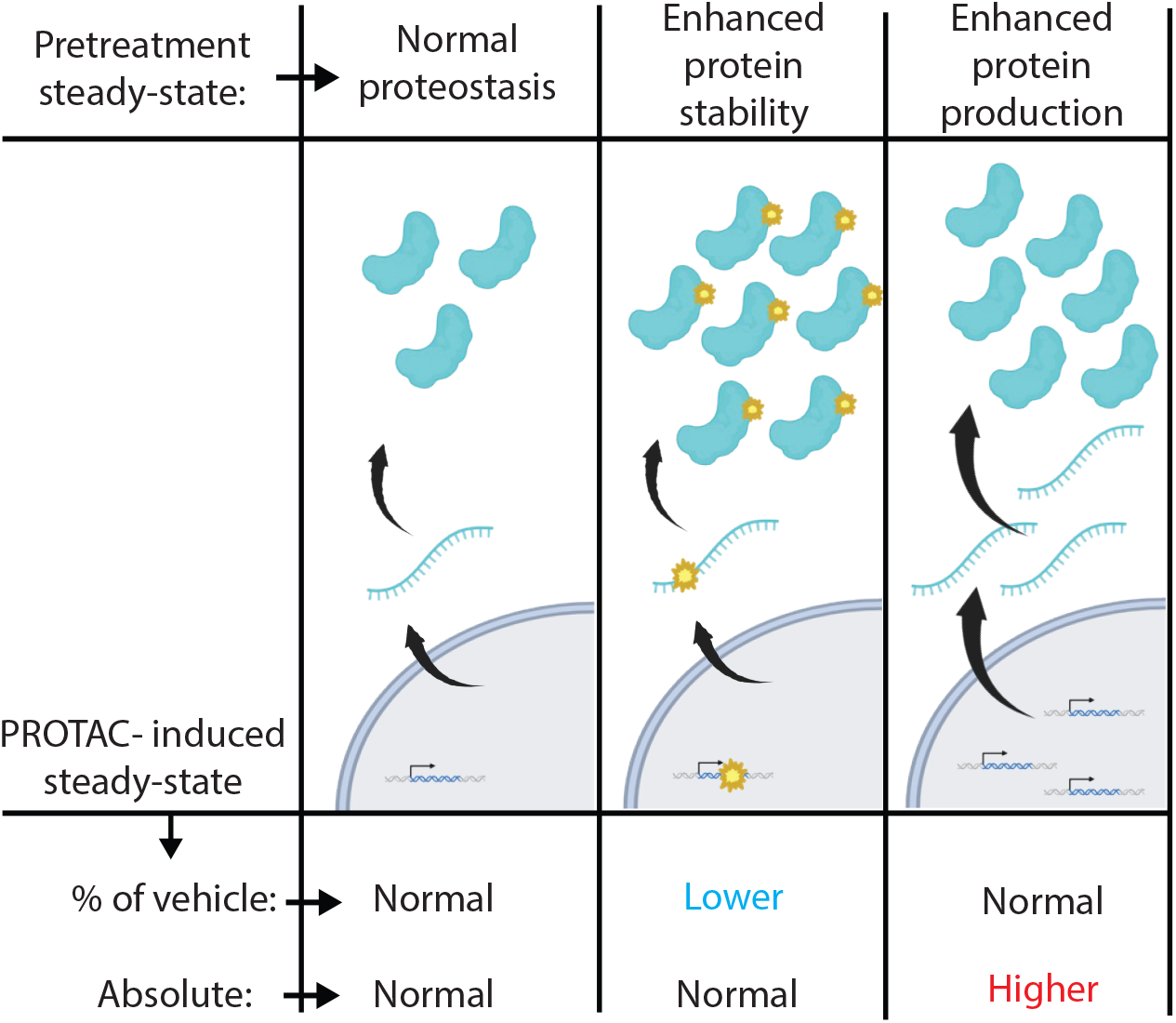
Model: differential effects of oncogene activation via protein stability versus transcriptional induction on PROTAC activity. Figure created using BioRender. Wood, A. (2025) https://BioRender.com/wv2rpyb

Our results demonstrate that oncogenic *CTNNB1* variants carrying mutations within the native degron increased steady-state protein levels up to 5-fold, consistent with their stabilising effects in human cancers (Figure 1, (Morin *et al*, 1997; Aberle *et al*, 1997)). Importantly, however, this elevated baseline abundance did not translate into increased resistance to PROTAC-induced degradation. Across three mutant alleles with graded effects on stability (Krishna *et al*, 2026), PROTAC treatment consistently converged *β*-catenin protein levels to similar absolute minima (Figure 2). Thus, although stabilising mutations raise the starting point from which degradation occurs, they do not alter the PROTAC-imposed steady-state. These results align with theoretical predictions that proteins with slow natural clearance are, in principle, highly susceptible to induced degradation (Bartlett & Gilbert, 2022; Vetma *et al*, 2024) and provide empirical support for the concept that induced degradation kinetics dominate over endogenous turnover. By contrast, oncogenic activation through increased transcriptional output imposed a different constraint on degrader activity. In a doxycycline-inducible system, raising the synthesis rate of *β*-catenin increased protein levels both before and after PROTAC treatment, leaving the fractional degradation maximum unchanged. This behaviour is consistent with a model in which PROTAC-mediated degradation establishes a new steady-state determined jointly by degradation rate and synthesis rate (Bartlett & Gilbert, 2022; Vetma *et al*, 2024). When synthesis is elevated, this steady-state shifts upward, limiting the depth of achievable depletion.

These observations have several implications for therapeutic development. First, preclinical degrader evaluation may benefit from incorporating models that vary both synthesis rate and stability, as oncogenic alleles encountered in the clinic reflect diverse proteostatic states. Second, tumours with high-level gene amplification may limit the achievable depth of degradation, providing a potential resistance pathway. This would be especially true in scenarios where gene copy number changes can evolve rapidly; for example, under conditions of chromosomal instability (Lukow *et al*, 2021), or when degrader target proteins are encoded on extrachromosomal circular DNA (Verhaak *et al*, 2019; Wong *et al*, 2026). While high level copy number gain of CTNNB1 is very rare (Figure 3A) it is far more common for other potential degrader targets (e.g. cMYC, nMYC, EGFR, (Zack *et al*, 2013)). Future work should investigate whether tumours adaptively increase synthesis rate in response to degrader pressure, analogous to cases of gene amplification observed during targeted kinase inhibitor treatment (Volpe *et al*, 2009; Corcoran *et al*, 2010). Third, because stabilising mutations do not limit the minimum achievable concentration under degrader treatment, patients with mutations that stabilise oncoproteins (e.g. missense mutations in CTNNB1 exon3 or APC frameshifts ((Korinek *et al*, 1997; Morin *et al*, 1997)) may remain highly responsive to degrader therapy despite elevated baseline protein levels.

Certain limitations of our work warrant discussion. We employed engineered degron tags and exogenous expression systems, which provided a reductionist system in which kinetic parameters associated with target protein turnover could be studied in isolation. We showed that the native *β*-catenin degron retained function in the tagged *CTNNB1* fusion and obtained similar results with VHL and CRBN-recruiting PROTACs. Although we see no logical reason why the core principles described in this work should differ across targets, or between tagged versus endogenous proteins, different substrate proteins and degrader ligands are expected to have distinct kinetic properties. Moreover, it is important to note that many other variables that were not examined in this study are likely to influence degrader activity across patients and clinical samples. Target protein proteostasis should therefore be viewed as one aspect in a much more complex picture.

In summary, we provide empirical evidence that distinct oncogenic activation mechanisms impose fundamentally different constraints on PROTAC activity. Stabilising mutations elevate steady-state protein levels but do not impede the minimal achievable abundance following degrader treatment, whereas transcriptional upregulation raises both pre- and post-treatment expression by increasing synthesis flux. These results offer a framework for interpreting degrader activity across diverse genetic backgrounds and underscore the importance of considering pre-treatment protein homeostasis in degrader drug development, resistance prediction, and personalised therapeutic strategies.

## Acknowledgments

The authors are grateful to Dr Kevin Myant for useful discussions, and to the Institute of Genetics and Cancer core facilities for technical support (Flow Cytometry and Advanced Imaging Resource). This work was funded by the Medical Research Council, UK (MC_PC_21040), and by a Medical Research Scotland PhD studentship (PhD-50695-2023).

## Declaration of Interests

AW has received sponsored lecture fees from Altos Labs, and is a paid consultant for Gemini Law. All other authors declare no competing financial interests.

## Declaration of generative AI and AI-assisted technologies

During the preparation of this work, the authors used Chat GPT in order to improve the readability of the text. After using this tool, the authors reviewed and edited the content as needed and take full responsibility for the content of the published article.

## Materials and Methods

### Generation and culture of stably expressing cell lines

The CTNNB1 degron reporter construct was generated by fusing the FKBP12^F36V^ (dTAG) and eGFP coding sequences to the C-terminus of the wildtype CTNNB1 open reading frame. Glycine-rich linker sequences ((GGGGS)3) separated CTNNB1 and dTAG, and dTAG and eGFP. These constructs were synthesised as clonal genes in pTwist Amp High Copy Cloning Vector by TWIST Bioscience, then gene fragments containing either S33F, T41A and S45F mutations in CTNNB1 were introduced by restriction cloning. Each full length ORF was then transferred into the pCDNA5-CAG-FRT expression vector (Ford *et al*., 2018), then transfected into Flp-In and/or T-REx HEK 293 cells (Invitrogen, R71407 and R78007) using Lipofectamine 3000. Transfections were performed using 2.5*µ*g of DNA, with a 9:1 ratio of the pcDNA vector and the Flp recombinase expression vector pOG44. Transfected cells were cultured in the presence of hygromycin B to select stable integrations, then GFP+ cells were purified by Fluorescence activated cell sorting (FACS). Integrated cells were used as pools in subsequent experiments, without clonal expansion. T-REx HeLa and Flp-In T-REx 293 cell lines were cultured in complete Dulbecco^*′*^s Modified Eagle^*′*^s Medium (DMEM) (Sigma-Aldrich, D5796) supplemented with 10% Fetal Calf Serum (FCS) (IGC Technical Services) and 1% Penicillin-Streptomycin solution (P 70 mg/l, S 130 mg/l, IGC Technical Services). All cells were incubated at 37°C, 5% CO2 and a humidified atmosphere. Cells were split and passaged once around 80% confluency was reached.

### Drug Treatments

For degrader dose response and time course experiments, 4 x 104 cells were seeded in individual wells of poly-L-lysine-coated 96 well plates (Sigma-Aldrich, P4707) 24 hours before treatment with dTAGV-1 and dTAG-13 then incubated at 37°C either for 2 hours at the indicated concentrations, or at 100nM for the indicated timepoints. To activate the Wnt pathway, cells were treated with GSK-3 *β* inhibitor (CHIR99021) for 24 h at a concentration of 10 nM. To activate transcription of CTNNB1 degron reporter transgenes, cells were incubated with 2.25*µ*M doxycycline for 24 hours before degrader treatment.

### Quantification of protein expression and degradation levels by flow cytometry

Cells were washed once with PBS then trypsinised to generate single cell suspensions for flow cytometry, and kept on ice until analysis. Data were collected using a CytoFLEX S (Beckman Coulter), then analysed using FlowJo 10. The corrected GFP fluorescence for each sample was determined by subtracting autofluorescence, measured in parallel from an equivalently-gated population of GFP negative cells. Then corrected fluorescence of the degrader-treated samples was normalised to 100% by dividing the corrected fluorescence of the sample by the corrected fluorescence of the vehicle-treated sample, multiplied by 100. Corrected fluorescence and normalised results were displayed as a dose-response curve, generated in R using the drc package (Ritz *et al*, 2015). The degradation maximum (D_max_) was determined by subtracting the percentage of remaining protein from 100%, selecting the highest percentage observed at any given concentration or timepoint. The half-maximal degradation concentration (DC_50_) was determined using the built-in model ED function in R for effective dose (ED) at 50% response. Boxplots were generated in R using either base R functions or, where required, functions from the qpcR package.

### Immunocytochemistry

Cells were plated on a coverslip in a 6-well plate a day prior. On the day, cells were washed with PBS containing calcium and magnesium (PBS++), followed by fixation with ice-cold 4% formaldehyde for 10 minutes. After fixation, cells underwent three washes with PBS++. Permeabilization was performed for 10 minutes in PBS containing 0.2% Triton X100, followed by additional washes. The cells were then blocked in 0.3% BSA in PBS for 1 hour and incubated with anti-tubulin and GFP-booster Alexa Fluor 488 (green, proteintech, gb2AF488) for an additional 1 hour. After the washing step, the coverslip was incubated with anti-rabbit Alexa Fluor 555 (red, Invitrogen, A31572) for 1 hour, followed by another round of washing before DAPI (blue) staining for 5 minutes. Finally, coverslips were mounted onto slides, and imaged using confocal microscopy. Image analysis was performed with CellProfiler, using segmentation techniques, where cytoplasm was set as a larger object and nucleus as smaller, which was extracted when quantifying the fluorescence of cytoplasm. This method enabled precise quantification of fluorescence across a minimum of 550 cells.

### qRT-PCR

Total RNA was isolated from cell pellets using TRIzol reagent, following the manufacturer’s protocol. Subsequently, the RNA underwent DNA removal using the DNA-Free kit (Invitrogen, AM1906) to eliminate any DNA contaminants. The RNA was then reverse transcribed into complementary DNA (cDNA) using the RevertAid H Minus First Strand cDNA Synthesis Kit (Thermo Fisher Scientific, K1632). qPCR primer pairs targeting Axin 2 (Forward: CAAACTTTCGCCAACCGTGGTTG, Reverse: GGTGCAAAGACATAGCCAGAACC) and *β*-Actin (Forward: CACCATTGGCAATGAGCGGTTC, Reverse: AGGTCTTTGCGGATGTCCACGT) (NM_004655) and *β*-Actin were employed for quantitative polymerase chain reaction (qPCR). SYBR Select Master Mix (Applied Biosystems, 4472908) protocol was utilized for qPCR, according to the manufacturer’s instructions, with amplification performed on the Bio-Rad CFX96 Touch instrument. The qPCR cycling conditions consisted of an initial activation step at 95^°^C for 5 min, followed by 39 cycles of denaturation at 95^°^C for 10 sec, annealing at 55^°^C for 10 sec, and extension at 72^°^C for 20 sec, with data acquisition at each step. A melting curve analysis was conducted with a temperature profile of 95^°^C for 10 sec, 65^°^C for 5 sec, and subsequent reduction by 0.5^°^C increments from 95^°^C. To evaluate the relative change in gene expression between cell lines expressing transgenic tagged *β*-catenin and those with wild-type *β*-catenin, the delta-delta Ct (ΔΔCt) method was employed. This method involved determining the difference in threshold cycle (Ct) values between the target gene (Axin 2) and the reference gene (*β*-Actin) for each sample. The Ct values of the target gene were then normalized to those of the reference gene to obtain the ΔCt. The (ΔΔCt) was then calculated by subtracting the ΔCt value of the control sample from the ΔCt value of each sample. Finally, the relative change in gene expression between the experimental and control conditions was determined using the formula 2^*−*ΔΔ*Ct*^.

### Modelling of degron reporter protein structure

3D protein models were generated using AlphaFold2 using the primary amino acid sequence of each construct to generate a PDB file. Models were displayed using ChimeraX.

### Analysis of gene copy number and mRNA expression in TCGA datasets

RNA-Seq (RSEM) and CTNNB1 gene copy number data (GISTIC2.0) were obtained from the TCGA Pan Cancer Atlas dataset, accessed via cBioportal. WNT-active tumours were identified among Liver Hepatocellular Carcinoma (LIHC), Endometrial Carcinoma (UCEC), Colorectal Adenocarcinoma (COAD) and Stomach Adenocarcinoma (STAD) datasets by filtering on samples with mutations in APC, CTNNB1 or RNF43.

---

**Figure S1:**
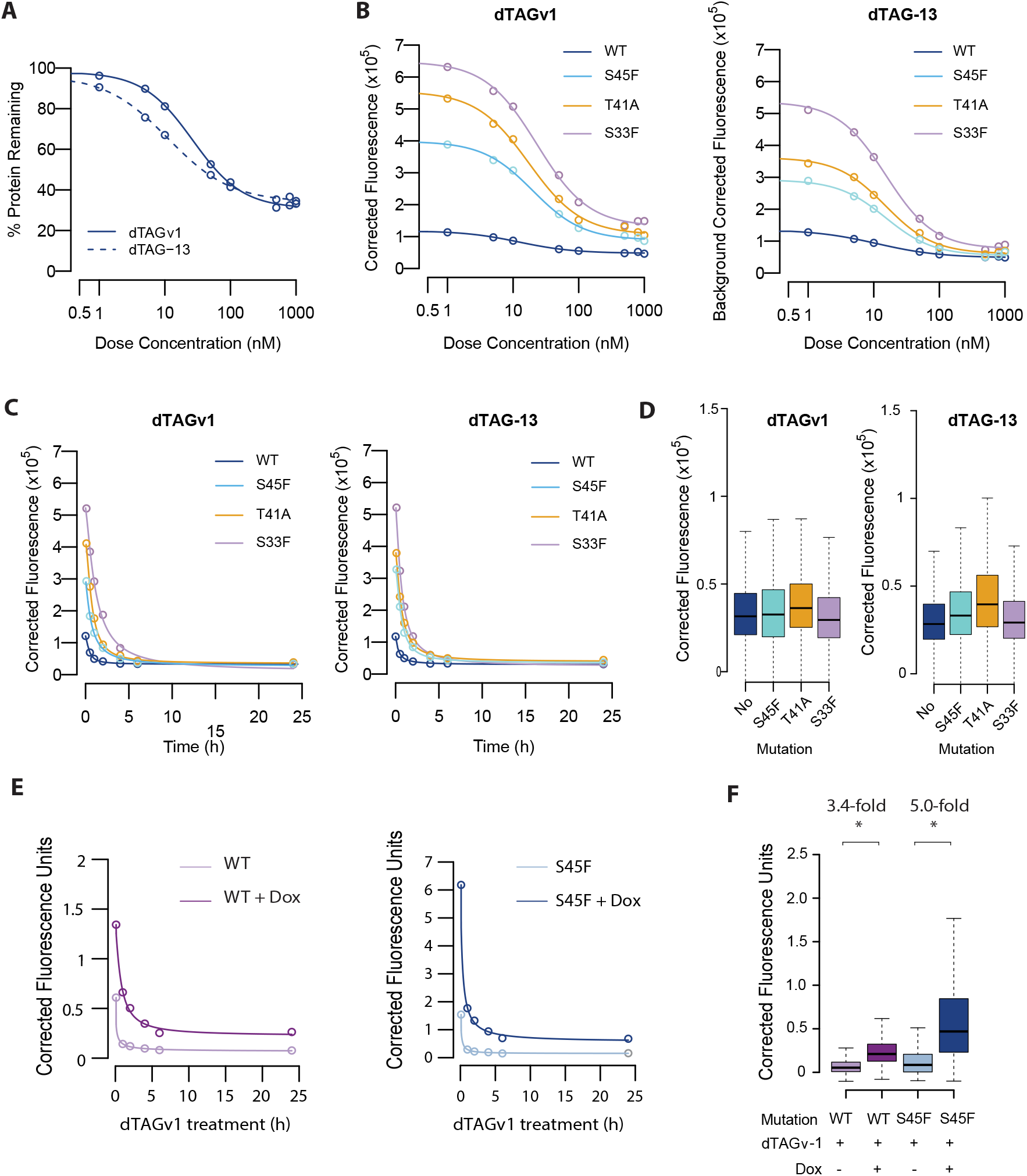
Second, independent biological replicates for degradation dose response and time course experiments in Figures 2 and 4. A. Dose response experiments shows degradation of the WT CTNNB1 reporter via dTAG-v1 and dTAG-13. Replicate experiment for Figure 1E. B. Dose response experiments for WT and mutant CTNNB1 reporters via dTAGv1 and dTAG-13. Replicate experiment for Figure 2A C. Time course experiments show degradation of WT and mutant CTNNB1 reporters via dTAGv1 and dTAG-13. Independent replicate experiment for Figure 2B. D. Boxplots show the distribution of GFP fluorescence levels mong cells (>1000) treated with dTAGv1 and dTAG-13 for 24 hours. Horizontal lines show the median fluorescence, boxes show the interquartile range and whiskers show the range. Independent replicate from Figure 2C. E. Time course experiment shows degradation of WT and S45 mutation reporters in the presence or absence of transcriptional induction via doxycycline. Independent replicate experiment for Figure 4C. F. Boxplots show the distribution of GFP fluorescence levels among cells (>1000) treated first with doxycycline for 24h, then with doxycycline and either dTAGv1 or dTAG-13 for a further 24 hours. Horizontal lines show the median fluorescence, boxes show the interquartile range and whiskers show the range. Independent replicate experiment for Figure 4D.

**Figure S2:**
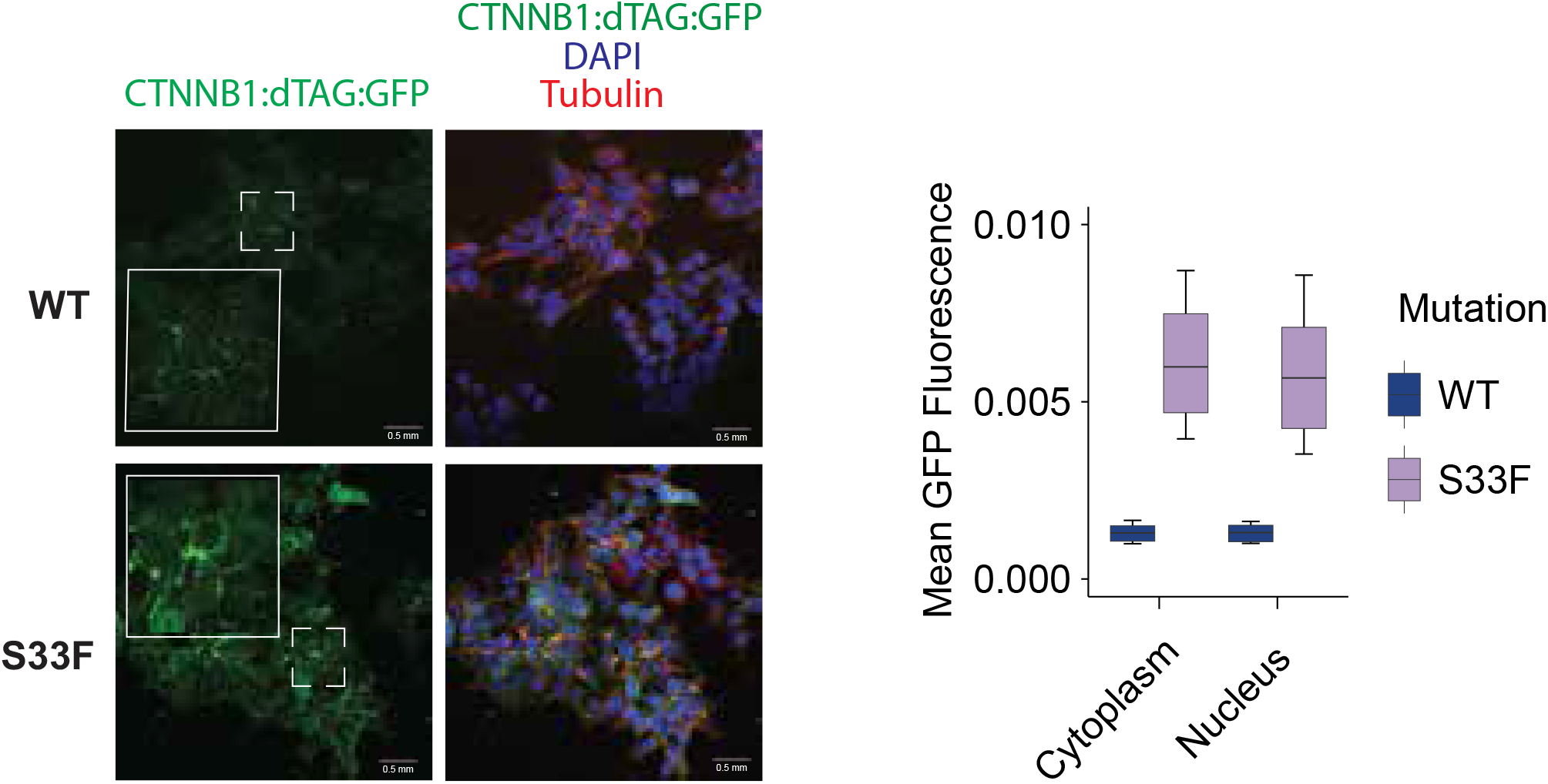
Native degron mutations increase nuclear levels of CTNNB1 degron reporter protein Immunofluorescence imaging was performed in transgene expressing HEK cells using anti-tubulin (red) and GFP booster (green) antibodies. Mean GFP pixel intensity was measured separately in the nucleus and cytoplasm of >500 single cells from one experiment. Horizontal lines show median values, boxes show the interquartile range, whiskers show the range.

## Notes

### Competing Interest Statement

The authors have declared no competing interest.

## References

Aberle H, Bauer A, Stappert J, Kispert A & Kemler R (1997) β-catenin is a target for the ubiquitin–proteasome pathway. EMBO J 16: 3797–3804

Ambrosi G, Voloshanenko O, Eckert AF, Kranz D, Nienhaus GU & Boutros M (2022) Allele-specific endogenous tagging and quantitative analysis of β-catenin in colorectal cancer cells. eLife 11: e64498

Bartlett DW & Gilbert AM (2022) Translational PK–PD for targeted protein degradation. Chem Soc Rev 51: 3477–3486

Bass AJ, Watanabe H, Mermel CH, Yu S, Perner S, Verhaak RG, Kim SY, Wardwell L, Tamayo P, Gat-Viks I, et al (2009) SOX2 is an amplified lineage-survival oncogene in lung and esophageal squamous cell carcinomas. Nat Genet 41: 1238–1242

Békés M, Langley DR & Crews CM (2022) PROTAC targeted protein degraders: the past is prologue. Nat Rev Drug Discov 21: 181–200

Boudreau MW, Freire VF, Corbett SC, Martínez-Fructuoso L, Kumar R, Thornburg CC, Akee RK, Peyser BD, Jiang Q, Splaine J, et al (2025) Positive Selection Screen for Natural Product β-Catenin Inactivators. bioRxiv: Prepr Serv Biol: 2025.08.27.671140

Chan JJ, Zhang B, Chew XH, Salhi A, Kwok ZH, Lim CY, Desi N, Subramaniam N, Siemens A, Kinanti T, et al (2022) Pan-cancer pervasive upregulation of 3 UTR splicing drives tumourigenesis. Nat Cell Biol 24: 928–939

Clevers H & Nusse R (2012) Wnt/β-Catenin Signaling and Disease. Cell 149: 1192–1205

Corcoran RB, Dias-Santagata D, Bergethon K, Iafrate AJ, Settleman J & Engelman JA (2010) BRAF Gene Amplification Can Promote Acquired Resistance to MEK Inhibitors in Cancer Cells Harboring the BRAF V600E Mutation. Sci Signal 3: ra84

Gao C, Wang Y, Broaddus R, Sun L, Xue F & Zhang W (2017) Exon 3 mutations of CTNNB1 drive tumorigenesis: a review. Oncotarget 9: 5492–5508

Gowans FA, Forte N, Hatcher J, Huang OW, Wang Y, Poblano BEA, Wertz IE & Nomura DK (2024) Covalent Degrader of the Oncogenic Transcription Factor β Catenin. J Am Chem Soc 146: 16856–16865

Hinterndorfer M, Spiteri VA, Ciulli A & Winter GE (2025) Targeted protein degradation for cancer therapy. Nat Rev Cancer 25: 493–516

Korinek V, Barker N, Morin PJ, Wichen D van, Weger R de, Kinzler KW, Vogelstein B & Clevers H (1997) Constitutive Transcriptional Activation by a β-Catenin-Tcf Complex in APC/ Colon Carcinoma. Science 275: 1784–1787

Krishna A, Meynert A, Dolt KS, Kelder M, Mesropian A, Ewing A, Brouwers C, Claassens JW, Linssen MM, Sheraz S, et al (2026) Mutational scanning reveals oncogenic CTNNB1 mutations have diverse effects on signaling. Nat Genet: 1–10

Liu J, Tokheim C, Lee JD, Gan W, North BJ, Liu XS, Pandolfi PP & Wei W (2021) Genetic fusions favor tumorigenesis through degron loss in oncogenes. Nat Commun 12: 6704

Lockwood WW, Chari R, Coe BP, Girard L, MacAulay C, Lam S, Gazdar AF, Minna JD & Lam WL (2008) DNA amplification is a ubiquitous mechanism of oncogene activation in lung and other cancers. Oncogene 27: 4615–4624

Lukow DA, Sausville EL, Suri P, Chunduri NK, Wieland A, Leu J, Smith JC, Girish V, Kumar AA, Kendall J, et al (2021) Chromosomal instability accelerates the evolution of resistance to anti-cancer therapies. Dev Cell 56: 2427-2439.e4

Mészáros B, Kumar M, Gibson TJ, Uyar B & Dosztányi Z (2017) Degrons in cancer. Sci Signal 10

Morin PJ, Sparks AB, Korinek V, Barker N, Clevers H, Vogelstein B & Kinzler KW (1997) Activation of β-Catenin-Tcf Signaling in Colon Cancer by Mutations in β-Catenin or APC. Science 275: 1787–1790

Nabet B, Ferguson FM, Seong BKA, Kuljanin M, Leggett AL, Mohardt ML, Robichaud A, Conway AS, Buckley DL, Mancias JD, et al (2020) Rapid and direct control of target protein levels with VHL-recruiting dTAG molecules. Nat Commun 11: 4687

Nabet B, Roberts JM, Buckley DL, Paulk J, Dastjerdi S, Yang A, Leggett AL, Erb MA, Lawlor MA, Souza A, et al (2018) The dTAG system for immediate and target-specific protein degradation. Nat Chem Biol 14: 431–441

Otto T, Horn S, Brockmann M, Eilers U, Schüttrumpf L, Popov N, Kenney AM, Schulte JH, Beijersbergen R, Christiansen H, et al (2009) Stabilization of N-Myc Is a Critical Function of Aurora A in Human Neuroblastoma. Cancer Cell 15: 67–78

Rebouissou S, Franconi A, Calderaro J, Letouzé E, Imbeaud S, Pilati C, Nault J, Couchy G, Laurent A, Balabaud C, et al (2016) Genotype-phenotype correlation of CTNNB1 mutations reveals different ß-catenin activity associated with liver tumor progression. Hepatology 64: 2047–2061

Riching KM, Mahan S, Corona CR, McDougall M, Vasta JD, Robers MB, Urh M & Daniels DL (2018) Quantitative Live-Cell Kinetic Degradation and Mechanistic Profiling of PROTAC Mode of Action. ACS Chem Biol 13: 2758–2770

Ritz C, Baty F, Streibig JC & Gerhard D (2015) Dose-Response Analysis Using R. PLoS ONE 10: e0146021

Sakamoto KM, Kim KB, Kumagai A, Mercurio F, Crews CM & Deshaies RJ (2001) Protacs: Chimeric molecules that target proteins to the Skp1–Cullin–F box complex for ubiquitination and degradation. Proc Natl Acad Sci 98: 8554–8559

Simonetta KR, Taygerly J, Boyle K, Basham SE, Padovani C, Lou Y, Cummins TJ, Yung SL, Soly SK von, Kayser F, et al (2019) Prospective discovery of small molecule enhancers of an E3 ligase-substrate interaction. Nat Commun 10: 1402

Taguchi K & Yamamoto M (2017) The KEAP1–NRF2 System in Cancer. Front Oncol 7: 85

Uniyal P, Kashyap VK, Behl T, Parashar D & Rawat R (2025) KRAS Mutations in Cancer: Understanding Signaling Pathways to Immune Regulation and the Potential of Immunotherapy. Cancers 17: 785

Verhaak RGW, Bafna V & Mischel PS (2019) Extrachromosomal oncogene amplification in tumour pathogenesis and evolution. Nat Rev Cancer 19: 283–288

Vetma V, Perez LC, Eliaš J, Stingu A, Kombara A, Gmaschitz T, Braun N, Ciftci T, Dahmann G, Diers E, et al (2024) Confounding Factors in Targeted Degradation of Short-Lived Proteins. ACS Chem Biol 19: 1484–1494

Volpe G, Panuzzo C, Ulisciani S & Cilloni D (2009) Imatinib resistance in CML. Cancer Lett 274: 1–9

Williams RD, Chagtai T, Alcaide-German M, Apps J, Wegert J, Popov S, Vujanic G, Tinteren H van, Heuvel-Eibrink MM van den, Kool M, et al (2015) Multiple mechanisms of MYCN dysregulation in Wilms tumour. Oncotarget 6: 7232–7243

Winter GE, Buckley DL, Paulk J, Roberts JM, Souza A, Dhe-Paganon S & Bradner JE (2015) Phthalimide conjugation as a strategy for in vivo target protein degradation. Science 348: 1376–1381

Wong IT-L, Yi H, Melillo B, Cravatt BF, Chang HY & Mischel PS (2026) Targeting extrachromosomal DNA in human cancers. Nat Rev Drug Discov: 1–16

Zack TI, Schumacher SE, Carter SL, Cherniack AD, Saksena G, Tabak B, Lawrence MS, Zhang C-Z, Wala J, Mermel CH, et al (2013) Pan-cancer patterns of somatic copy number alteration. Nat Genet 45: 1134–1140

Zengerle M, Chan K-H & Ciulli A (2015) Selective Small Molecule Induced Degradation of the BET Bromodomain Protein BRD4. ACS Chem Biol 10: 1770–1777

